# The Omicron variant BA.1.1 presents a lower pathogenicity than B.1 D614G and Delta variants in a feline model of SARS-CoV-2 infection

**DOI:** 10.1101/2022.06.15.496220

**Authors:** Mathias Martins, Gabriela M. do Nascimento, Mohammed Nooruzzaman, Fangfeng Yuan, Chi Chen, Leonardo C. Caserta, Andrew D. Miller, Gary R. Whittaker, Ying Fang, Diego G. Diel

## Abstract

Omicron (B.1.1.529) is the most recent SARS-CoV-2 variant of concern (VOC), which emerged in late 2021 and rapidly achieved global predominance in early 2022. In this study, we compared the infection dynamics, tissue tropism and pathogenesis and pathogenicity of SARS-CoV-2 D614G (B.1), Delta (B.1.617.2) and Omicron BA.1.1 sublineage (B.1.1.529) variants in a highly susceptible feline model of infection. While D614G- and Delta-inoculated cats became lethargic, and showed increased body temperatures between days 1 and 3 post-infection (pi), Omicron-inoculated cats remained subclinical and, similar to control animals, gained weight throughout the 14-day experimental period. Intranasal inoculation of cats with D614G- and the Delta variants resulted in high infectious virus shedding in nasal secretions (up to 6.3 log10 TCID_50_.ml^-1^), whereas strikingly lower level of viruses shedding (<3.1 log10 TCID_50_.ml^-1^) was observed in Omicron-inoculated animals. In addition, tissue distribution of the Omicron variant was markedly reduced in comparison to the D614G and Delta variants, as evidenced by *in situ* viral RNA detection, *in situ* immunofluorescence, and quantification of viral loads in tissues on days 3, 5, and 14 pi. Nasal turbinate, trachea, and lung were the main - but not the only - sites of replication for all three viral variants. However, only scarce virus staining and lower viral titers suggest lower levels of viral replication in tissues from Omicron-infected animals. Notably, while D614G- and Delta-inoculated cats had severe pneumonia, histologic examination of the lungs from Omicron-infected cats revealed mild to modest inflammation. Together, these results demonstrate that the Omicron variant BA.1.1 is less pathogenic than D614G and Delta variants in a highly susceptible feline model.

**Author Summary:** The SARS-CoV-2 Omicron (B.1.1.529) variant of concern (VOC) emerged in South Africa late in 2021 and rapidly spread across the world causing a significant increase in the number of infections. Importantly, this variant was also associated with an increased risk of reinfections. However, the number of hospitalizations and deaths due to COVID-19 did not follow the same trends. These early observations, suggested effective protection conferred by immunizations and/or overall lower virulence of the highly mutated variant virus. In this study we present novel evidence demonstrating that the Omicron BA.1.1 variant of concern (VOC) presents a lower pathogenicity when compared to D614G- or Delta variants in cats. Clinical, virological and pathological evaluations revealed lower disease severity, viral replication and lung pathology in Omicron-infected cats when compared to D614G and Delta variant inoculated animals, confirming that Omicron BA.1.1 is less pathogenic in a highly susceptible feline model of infection.

## Introduction

More than two years after the first reported infections with severe acute respiratory syndrome coronavirus 2 (SARS-CoV-2) in a cluster of people in Wuhan, Hubei province in China, the virus remains circulating and evolving in the human population worldwide. The SARS-CoV-2 is a positive sense, single-stranded RNA virus that belongs to the *Sarbecovirus* subgenus, within the *Betacoronavirus* genus, of the family *Coronaviridae* [1]. The virus is closely related to other sarbecoviruses identified in horseshoe bats in China, which are currently considered the most likely source of the ancestral virus from which SARS-CoV-2 originated before it spillover into humans, potentially via an yet unidentified intermediate animal host [2,3]. Despite the implementation of major public health measures in an attempt to prevent the spread of SARS-CoV-2, and the rapid development of effective vaccines and antivirals against severe coronavirus disease 2019 (COVID-19), the pandemic continues to evolve with the emergence of new SARS-CoV-2 variants causing significant new waves of infection worldwide.

The first described mutation on the SARS-CoV-2 genome was the D614G substitution in the Spike (S) protein, which was detected in February 2020 and linked to increased transmissibility of new B.1 variant viruses [4], which quickly became the predominant SARS-CoV-2 lineage circulating worldwide. The next and more significantly diverse SARS-CoV-2 variants to emerge were B.1.1.7 (alpha) in the United Kingdom, B.1.351 (beta) in South Africa and P.1 (gamma) in Brazil. These variants retained the D614G S gene mutation with each of them presenting a characteristic constellation of mutations totaling 11, 10 and 12 amino acid substitutions only in the S gene, respectively. Some of these mutations resulted in escape from antibody therapy, enhance binding to the human ACE2 receptor and increased transmissibility in humans, which led to the classification of these viruses as variants of concern (VOC) by the World Health Organization (WHO) [5–7]. The SARS-CoV-2 lineage B.1.617.2 (Delta), containing 10 S gene mutations, was described for the first time in late 2020 in India, and by mid-2021, it became the predominant VOC circulating worldwide with the virus being described in more than 180 countries [7–9]. In addition to increased transmissibility, the Delta variant was also associated with vaccine breakthrough infections and increased pathogenicity in humans and animal models [10,11]. The most recent SARS-CoV-2 VOC B.1.1.529 (Omicron), presenting 37 mutations in the S gene, emerged in late 2021 in South Africa, and rapidly achieved global predominance in early 2022 [12]. Although the initial Omicron sublineages BA.1 and BA.1.1 were able to spread with unprecedented speed across the globe, causing a significant surge in the number of cases, the number of hospitalizations and deaths due to COVID-19 did not follow the same trend, suggesting effectiveness of vaccine induced immunity following booster shots or perhaps lower pathogenicity of the newly emergent variant. Notably, several studies demonstrated that immunity elicited by booster immunizations is required for effective neutralization of the Omicron variant [13–15]. Additionally, experimental studies show an overall lower pathogenicity of Omicron BA.1 and BA.1.1 lineages in hACE2 transgenic mice and hamster models for SARS-CoV-2 infection [14–18].

A better understanding of the infectivity and pathogenesis of SARS-CoV-2 VOCs is critical for development of improved vaccines and therapeutics to effectively control the COVID-19 pandemic. In this study, we characterized/compared the infection dynamics, tissue tropism and pathogenicity of SARS-CoV-2 D614G (B.1), Delta (B.1.617.2) and Omicron BA.1.1 (B.1.1.529) variants in a highly susceptible feline model.

## Results

### SARS-CoV-2 Omicron BA.1.1 leads to subclinical infection and limited viral shedding in cats

The dynamics of infection, virus replication, and viral shedding of SARS-CoV-2 D614G, Delta and Omicron variants were assessed in adult cats (Fig 1A). The animals were housed individually in Horsfall HEPA filter cages in the animal biosafety level 3 (ABSL-3) facility at Cornell University throughout the 14-day experimental period. Following inoculation, clinical parameters, including rectal temperature, body weight, and clinical signs of respiratory disease, were monitored daily. While D614G- and Delta-inoculated cats became lethargic, and showed significantly increased body temperatures between days 1 and 3 post-infection (pi) (Fig 1B), Omicron-inoculated cats remained subclinical. Additionally, D614G- and Delta-inoculated cats lost or maintained steady body weights throughout the 14-day experimental period, while Omicron-inoculated cats gained weight (3-9% of their initial body weight) similar to what was observed in the control mock-inoculated animals (Fig 1C).

**Figure 1.**
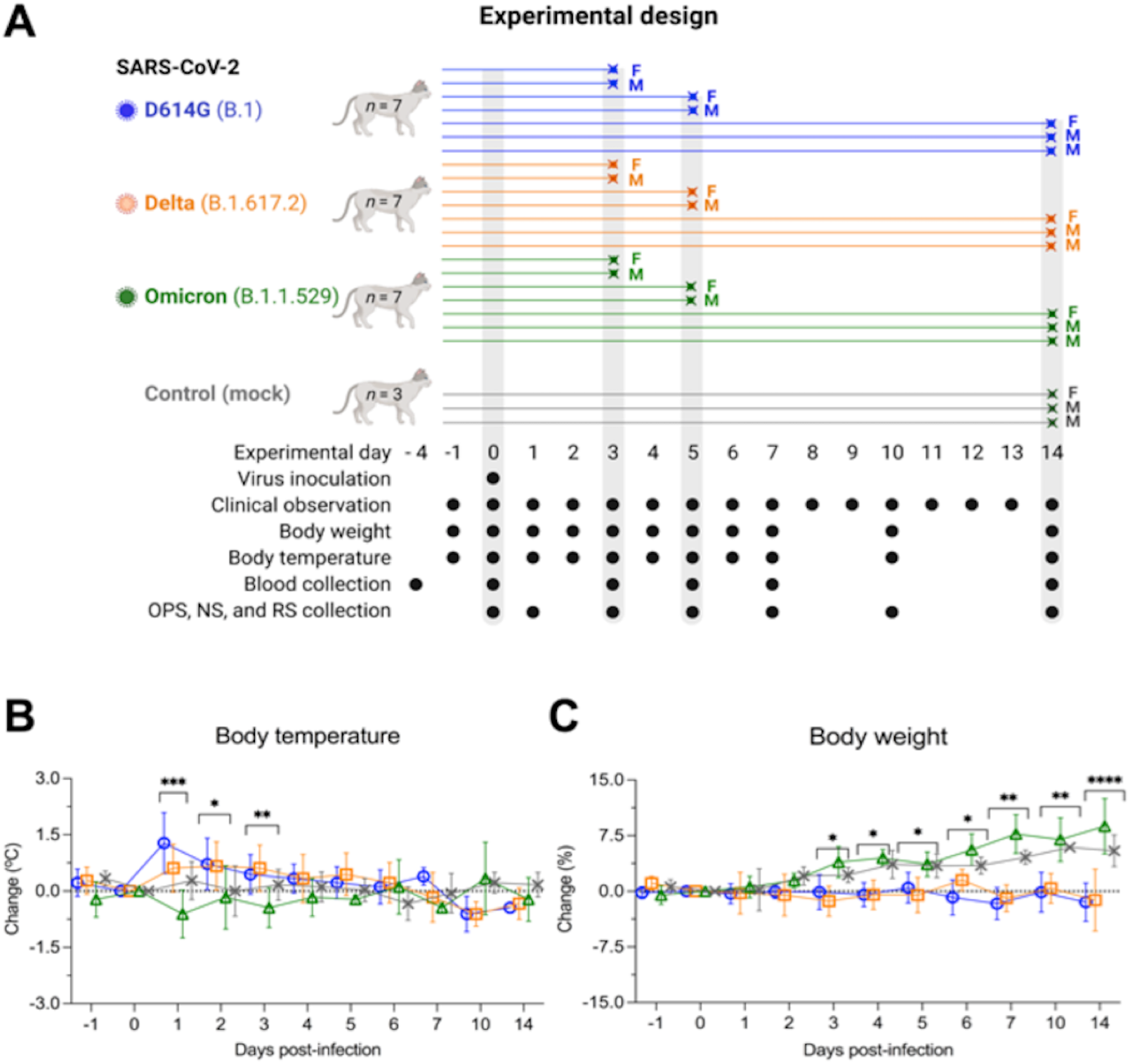
Experimental design, body temperature and weight changes following SARS-CoV-2 inoculation in cats. Schematic representation of the experimental design of infection study in adult cats (24-40-month-old) males and females. Animals were allocated in three inoculated groups (*n* = 7 per group) and a control group (mock inoculated) (*n* = 3). Animals were inoculated intranasally with 1 ml (0.5 ml per nostril) of virus suspension containing 5 x 10^5^ PFU of SARS-CoV-2 D614G (B.1), Delta (B.1.617.2), or Omicron BA.1.1 (B.1.1.529), or 1 ml of cell culture supernatant media (control mock inoculated). On day 3 and 5 post-infection (pi), two cats from each group (one female and one male) were humanely euthanized and the remaining cats (one female and two males) were maintained until day 14 pi. Respiratory secretion, feces and serum were collected on the specific days as indicated (**A)**. NS = nasal swab; OPS = oropharyngeal swab; RS = rectal swab; F = female; M = male. Body temperature (**B**) and body weight changes (**C**) following intranasal SARS-CoV-2 D614G (B.1), Delta (B.1.617.2), or Omicron BA.1.1 (B.1.1.529) inoculation throughout the 14-day experimental period. Data are presented as means ± standard deviation. * *p <* 0.05; ** *p <* 0.01; *** *p <* 0.005; **** *p <* 0.0001.

To assess virus replication and shedding dynamics of SARS-CoV-2 following inoculation, nasal and oropharyngeal secretions and feces were collected using nasal (NS), oropharyngeal (OPS), and rectal swabs (RS) (Fig 1A). The samples were initially tested for the presence of SARS-CoV-2 RNA by real-time reverse transcriptase PCR (rRT-PCR). Viral RNA was detected between days 1 and 14 pi in nasal and oropharyngeal secretions in the inoculated cats, regardless of the virus variant used (Fig 2A and B), with higher viral RNA loads detected between days 3 and 5 pi, which decreased thereafter through day 14 pi (Fig 2A and B). Viral RNA load was higher in nasal secretions of D614G- and Delta-inoculated animals when compared to Omicron-inoculated cats throughout the experiment (*p* < 0.001). Viral RNA load in oropharyngeal secretions was significantly lower in the Omicron-inoculated cats than in D614G- and Delta-inoculated animals on days 1 and 10 pi (*p* < 0.001) (Fig 2B). Shedding of viral RNA in feces was markedly lower than in respiratory and oropharyngeal secretions and was characterized by intermittent detection of low amounts of viral RNA, with Omicron-inoculated animals presenting lower RNA levels in feces between days 3 and 10 pi (*p* < 0.01) (Fig 2C). All control cats (mock inoculated) remained negative through the 14-day experimental period.

**Figure 2.**
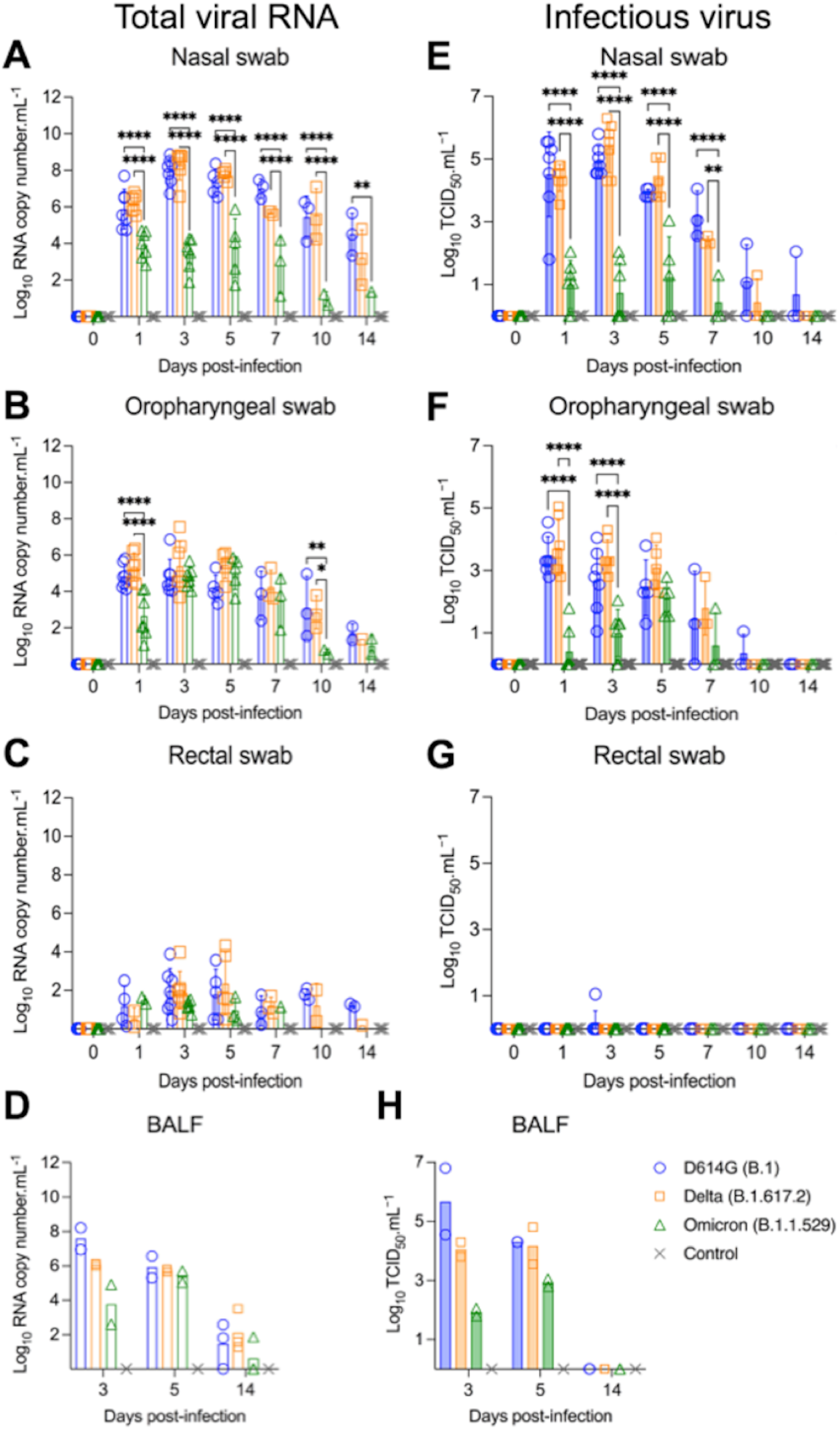
Viral shedding dynamics in respiratory secretions and feces following SARS-CoV-2 D614G, Delta, and Omicron BA.1.1 inoculation in cats. SARS-CoV-2 RNA load assessed in nasal (**A**), oropharyngeal secretions (**B**), and feces (**C**), collected on days 0, 1, 3, 5, 7, 10, and 14 post-infection (pi), and in bronchoalveolar lavage fluid (BALF) collected on 3, 5, and 14 pi (**D**). Samples were tested for the presence of SARS-CoV-2 RNA by real-time reverse transcriptase PCR (rRT-PCR) (genomic and subgenomic viral RNA load). Infectious SARS-CoV-2 load in nasal (**E**) and oropharyngeal secretions (**F**), feces (**G**), and BALF (**H)** assessed by virus titration in rRT-PCR-positive samples. Virus titers were determined using end point dilutions and expressed as TCID_50_.ml^-1^. Data are presented as means ± standard deviation. * *p <* 0.05; ** *p <* 0.01; *** *p <* 0.005; **** *p <* 0.0001.

The dynamics of infectious SARS-CoV-2 shedding was also assessed in nasal and oropharyngeal secretions and feces. All samples positive by rRT-PCR were tested for presence of infectious virus. High viral loads were detected in nasal and oropharyngeal secretions of D614G- and Delta-inoculated cats (Fig 2E and F). All D614G- and Delta-inoculated cats shed infectious SARS-CoV-2 between days 1–7 pi in the nasal secretions, with viral titers ranging from 2.3 to 6.3 log10 TCID_50_.ml^-1^, whereas Omicron-inoculated animals shed significantly lower viral titers (*p* < 0.001) ranging from 1.0 to 3.0 log10 TCID_50_.ml^-1^ (Fig 2E). Two out of three D614G-infected and one out of three Delta-infected cats shed infectious virus in nasal secretions on day 10 pi, and one D614G-inoculated cat on day 14 pi (Fig 2E). Cats inoculated with D614G and Delta viruses shed infectious SARS-CoV-2 between days 1–7 pi in the oropharyngeal secretions, with viral titers ranging from 1.3 to 5.0 log10 TCID_50_.ml^-1^, while Omicron-inoculated cats presented lower viral titers ranging from 1.0 to 2.8 log10 TCID_50_.ml^-1^ (Fig 2F). Animals inoculated with D614G- and Delta shed viral titers ranging from 2.8 to 5.0 log10 TCID_50_.ml^-1^ on days 1 and 3 pi in oral secretions, while Omicron-inoculated cats shed significantly lower viral titers (*p* < 0.001) (ranging from 1.0 to 2.0 log10 TCID_50_.ml^-1^) (Fig 2F). No significant infectious virus shedding was detected in feces in any of the groups (Fig. 2G).

We also assessed the viral RNA load and infectious virus titers in bronchoalveolar lavage fluid (BALF). Two cats from each SARS-CoV-2 inoculated group (D614G, Delta and Omicron) were humanely euthanized on days 3 and 5 pi, and three cats per group, including controls (mock inoculated), were euthanized on day 14 pi (Fig 1A). Viral RNA was detected in all inoculated animals on day 3 and 5 pi, regardless the inoculated virus (Fig 2D). All control cats (mock inoculated) tested negative by rRT-PCR. Infectious virus titers in BALF of cats infected with D614G or Delta varied from 3.8 to 6.8 log10 TCID_50_.ml^-1^, whereas in Omicron-infected animals, infectious virus titers ranged from 1.8 and 2.0 log10 TCID_50_.ml^-1^ on day 3 pi (Fig 2H). On day 5 pi, viral titers in cats infected with D614G or Delta varied from 3.8 to 4.8 log10 TCID_50_.ml^-1^, while infectious virus titers in BALF from Omicron-infected cats were 2.8 and 3.0 log10 TCID_50_.ml^-1^ (Fig 2H). Viral RNA was detected in BALF of 2/3, 3/3, and 1/3 cats of D614G, Delta, and Omicron-infected cats, respectively, on day 14 pi (Fig 2D). Regardless the inoculated virus, no infectious virus was detected in BALF on day 14 pi (Fig 2H). Together, these results demonstrate that SARS-CoV-2 Omicron BA.1.1 presents a lower pathogenicity when compared to the SARS-CoV-2 D614G and Delta variants in our feline model.

### SARS-CoV-2 Omicron BA.1.1 presents reduced replication in tissues

The tissue tropism and replication sites of SARS-CoV-2 D614G, Delta and Omicron variants were assessed following intranasal inoculation in cats. For this, nasal turbinate, palate/tonsil, retropharyngeal lymph node, trachea, lung, mediastinal lymph node, heart, liver, spleen, kidney, small intestine, and mesenteric lymph node were collected on days 3, 5 and 14 pi following euthanasia and processed for rRT-PCR, virus isolation, titrations and *in situ* hybridization and immunofluorescence staining. SARS-CoV-2 RNA was detected in several tissues sampled from each group, with higher viral RNA loads being detected on days 3 and 5 pi when compared to day 14 pi (Fig 3A-C). The highest total viral RNA loads were detected in the nasal turbinate on day 3 and 5 pi in D614G- and Delta-inoculated animals, with lower viral RNA loads detected in tissues from Omicron-inoculated cats (Fig 3A and B). To assess virus replication in tissues, subgenomic viral RNA was determined by qRT-PCR targeting the E gene [19]. Subgenomic viral RNA was consistently detected in nasal turbinate, palate/tonsil, trachea, retropharyngeal lymph node, and lung, from D614G- and Delta-inoculated cats on days 3 and 5 pi (Fig 3D and E). The highest subgenomic RNA loads were observed in nasal turbinate on days 3 and 5 pi, regardless the inoculated virus (Fig 3D and E). However, subgenomic RNA was detected only in nasal turbinate and trachea from Omicron-inoculated cats (Fig 3D and E). The subgenomic viral RNA loads detected in tissues from Omicron-inoculated cats was markedly lower than those detected in D614G- and Delta-inoculated animals (Fig 3D and E). On day 14 pi, viral RNA loads decreased when compared to early time points and subgenomic RNA was only detected in nasal turbinate, palate/tonsil, and retropharyngeal lymph nodes of D614G- and Delta-infected cats, whereas all tissues from the Omicron group tested negative (Fig 3F). All tissues from the control cats (mock inoculated) tested negative for total and subgenomic viral RNA by RT-PCR (Fig 3C and F).

**Figure 3.**
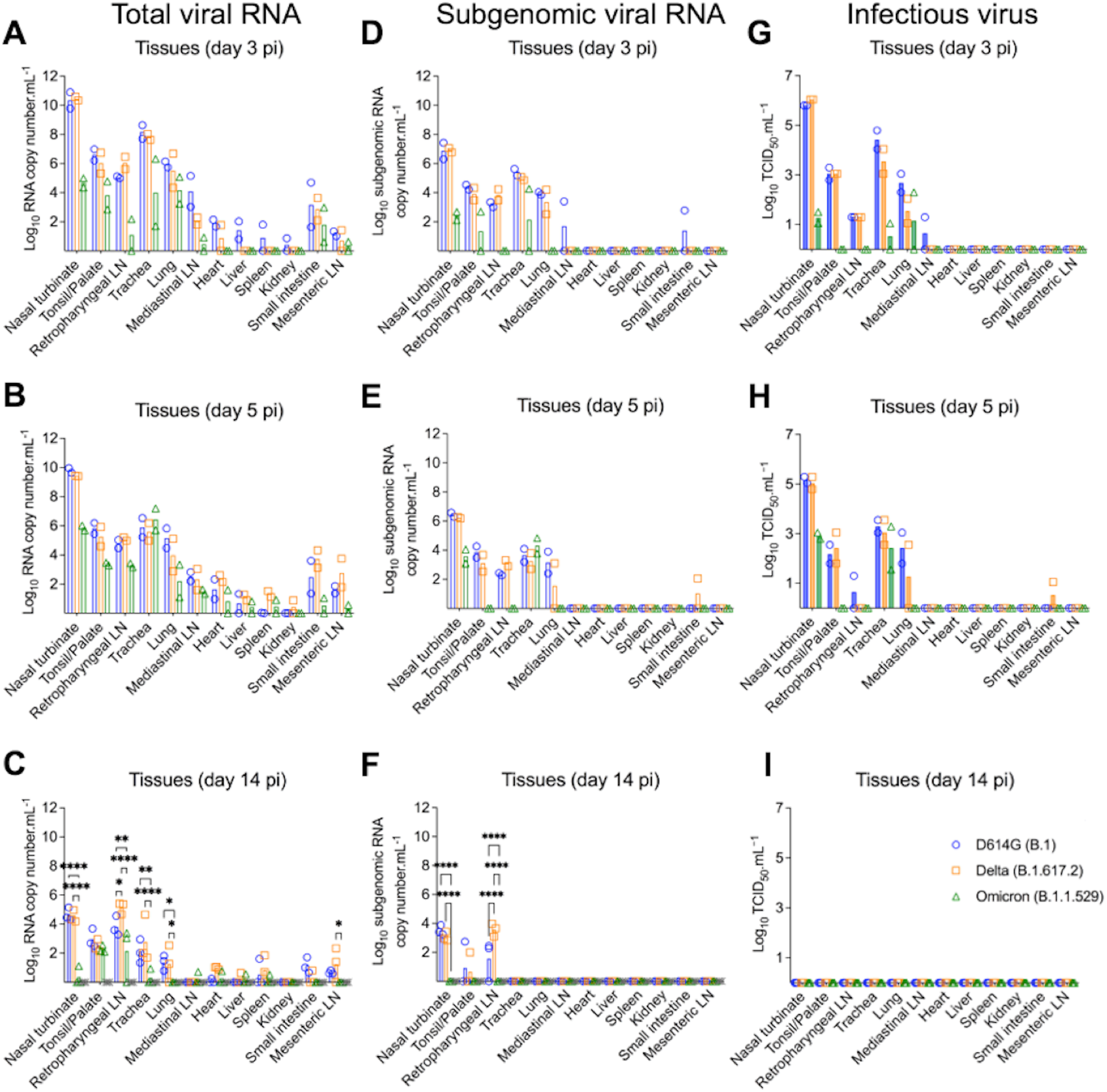
Tissue distribution of SARS-CoV-2 D614G, Delta, and Omicron BA.1.1 following virus inoculation in cats. Tissues were collected and processed for real-time reverse transcriptase PCR (rRT-PCR) and infectious virus titrations. Tissue distribution of SARS-CoV-2 RNA assessed by rRT-PCR. Samples from two cats per group were tested at day 3 (**A**) and 5 (**B**) post-infection (pi), and samples from three cats per group were tested at day 14 pi (**C**). Tissue distribution of subgenomic SARS-CoV-2 RNA (replicating RNA) assessed in tissue samples collected on days 3 (**D**), 5 (**E**), and 14 pi (**F**). Infectious SARS-CoV-2 in tissues assessed by virus titrations in rRT-PCR-positive samples obtained on days 3 (**G**), 5 (**H**) and 14 pi (**F**). Virus titers were determined using end point dilutions and expressed as TCID_50_.ml^-1^. Data are presented as means ± standard deviation. * *p <* 0.05; ** *p <* 0.01; *** *p <* 0.005; **** *p <* 0.0001.

The presence and amount of infectious SARS-CoV-2 in tissues was assessed by virus titrations in rRT-PCR positive tissues. Notably, detection of infectious virus and infectious viral loads were consistent with subgenomic viral RNA detection (Fig 3D-I). Infectious virus was consistently detected in a broad range of tissues on days 3 and 5 pi including nasal turbinate, palate/tonsil, retropharyngeal lymph node, trachea, and lung from D614G- and Delta-infected cats, whereas only nasal turbinate, trachea and lung from Omicron-infected cats were positive (Fig 3G an H). The highest viral titers were observed in the nasal turbinate (titers ranging 5.8 to 6.0 log10 TCID_50_.ml^-1^) from D614G- and Delta inoculated cats on days 3 and 5 pi, while markedly lower viral loads (1.0 to 3.0 log10 TCID_50_.ml^-1^) were detected in Omicron inoculated cats (Fig 3G-H). Interestingly, although on day 5 pi infectious virus titers detected in D614G- and Delta-inoculated animals were slightly lower than in day 3 pi, viral titers in tissues from Omicron-inoculated cats were slightly higher than viral titers detected on day 3 pi (Fig 3G and H). Together these results indicate reduced replication of the Omicron BA.1.1 virus in the feline model.

### SARS-CoV-2 Omicron BA.1.1 presents reduced replication in the respiratory tract of cats

The tissue distribution of SARS-CoV-2 in the respiratory tract and associated lymphoid tissues was assessed by *in situ* hybridization (ISH) and immunofluorescence (IFA) staining. For this, nasal turbinate, palate/tonsil, retropharyngeal lymph nodes, trachea, lung, and heart from virus inoculated animals were examined by ISH using the RNAscope^®^ ZZ technology and by IFA using a SARS-CoV-2-nucleoprotein (N) specific antibody. Viral RNA and the N protein were consistently detected in nasal turbinate, trachea, and lung from cats regardless the virus inoculated (Fig 4A and 5A). While intense labeling for the viral RNA and staining for the viral N protein were observed in tissues from D614G- and Delta-inoculated cats, only modest and sporadic staining was detected in tissues from Omicron-inoculated animals on days 3 and 5 pi (Fig 4A and 5A). The most abundant detection signals for viral RNA and N protein were observed in the nasal turbinate from D614G- and Delta-inoculated cats on days 3 and 5 pi (Fig 4A and 5A). When we compared the intensity and distribution of both viral RNA and N in tissues from the three inoculated groups (D614G, Delta and Omicron), the lower infectivity of the Omicron variant was evident by lower levels of viral RNA and N detection across all tissues tested (Fig 4A and 5A).

**Figure 4.**
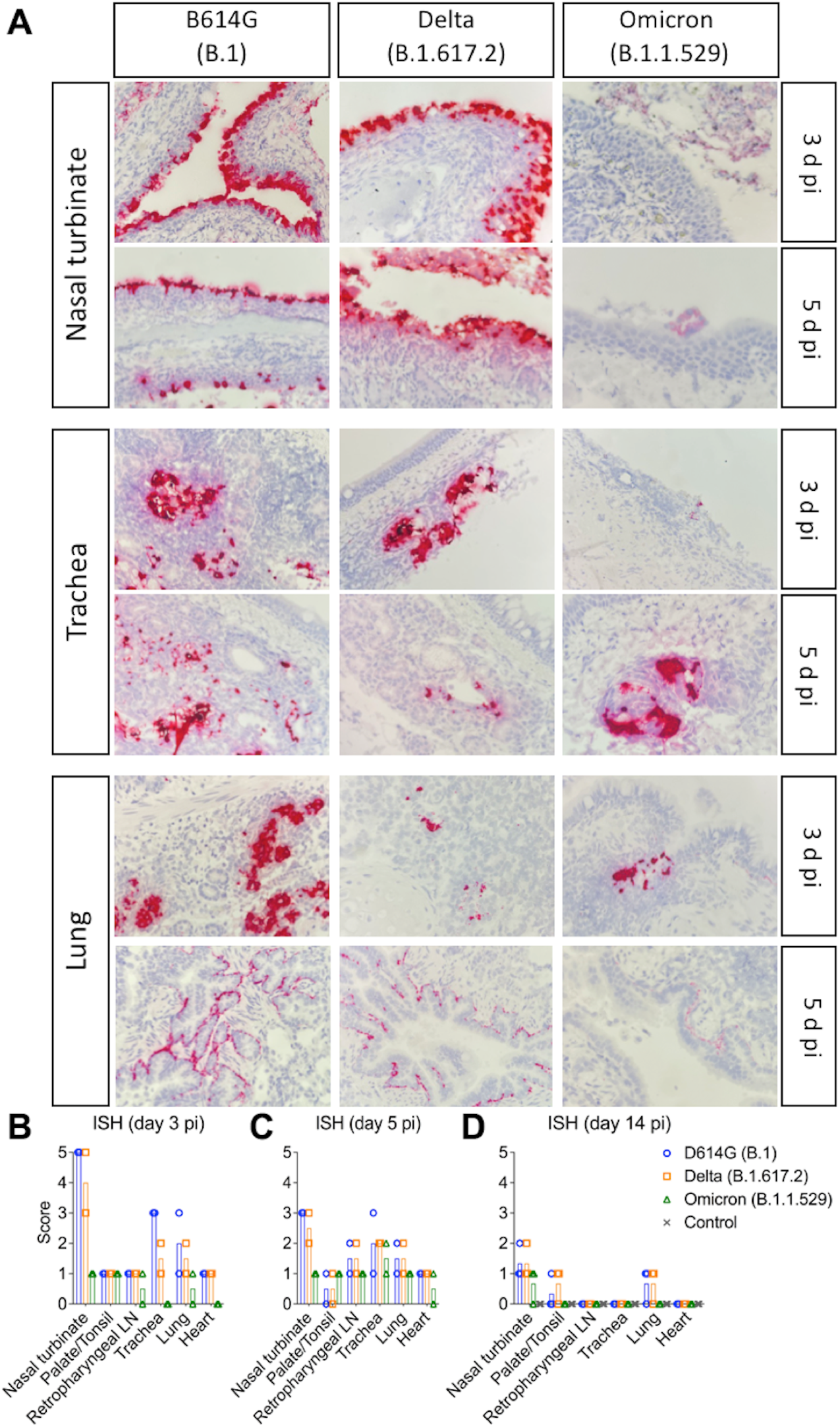
*In situ* hybridization (ISH) in tissues of SARS-CoV-2 D614G, Delta, and Omicron BA.1.1 inoculated cats. Nasal turbinate, trachea and lung samples collected from SARS-CoV-2 D614G (B.1), Delta (B.1.617.2), and Omicron (B.1.1.529) inoculated cats on days 3 and 5 post-infection (pi). Note, intensive labeling of viral RNA (SARS-CoV-2-Spike) (Red) on the tissues from D614G- and Delta-inoculated cats, while less abundant labeling on tissues from Omicron-infected animals on days 3 and 5 pi (**A**). In addition to above described tissues, palate/tonsil, retropharyngeal lymph node, and heart were evaluated. ISH scores (established as described in the Materials and Methods) for the tissues evaluated on days 3, 5, and 14 pi are presented in (**B**), (**C**), and (**D**), respectively.

**Figure 5.**
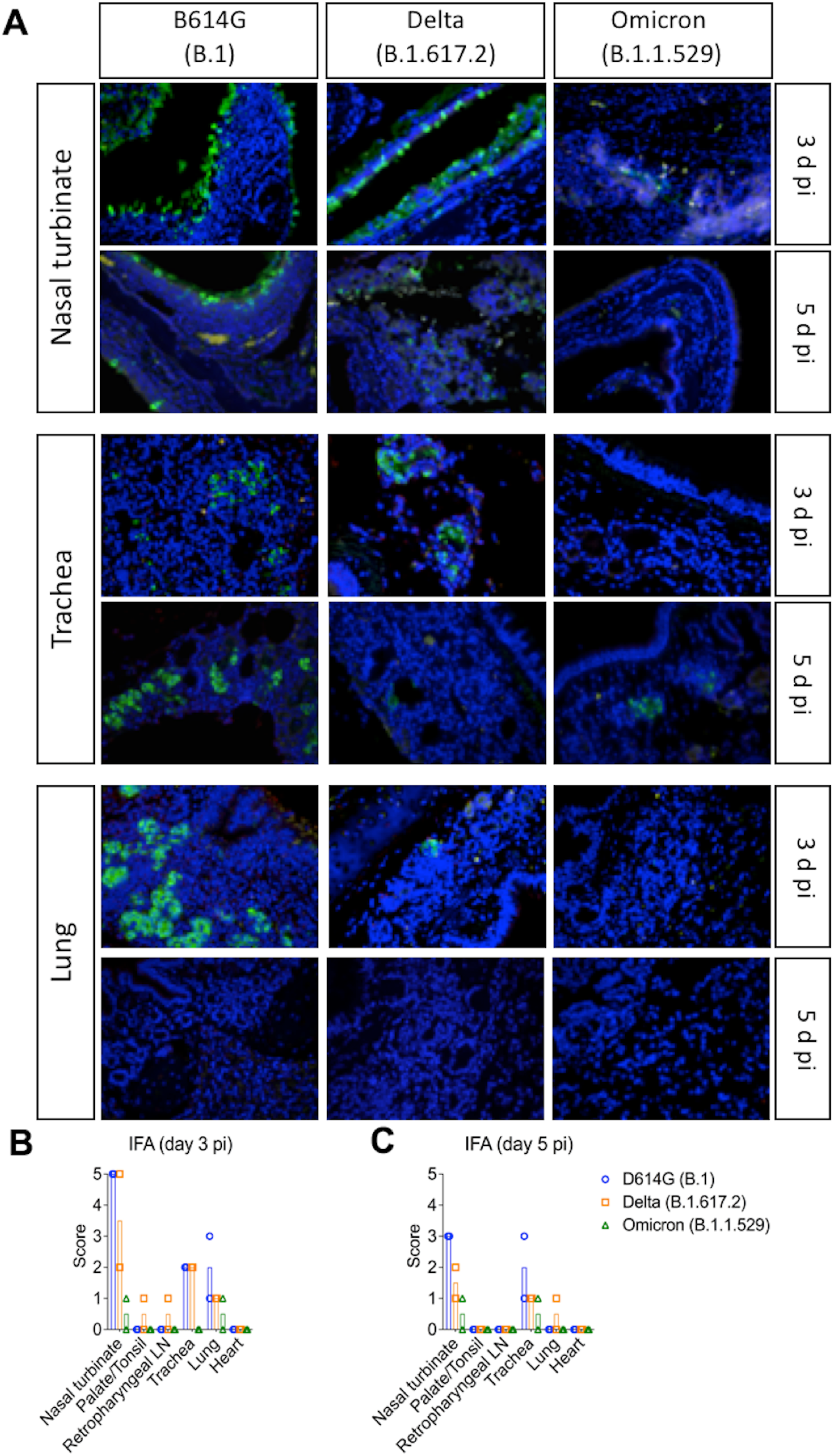
*In situ* immunofluorescence (IFA) in tissues of SARS-CoV-2 D614G, Delta, and Omicron BA.1.1 inoculated cats. Tissues were subjected to IFA using a monoclonal antibody against nucleocapsid protein (NP) of SARS-CoV-2. Nasal turbinate, trachea and lung were collected from SARS-CoV-2 D614G (B.1), Delta (B.1.617.2), and Omicron (B.1.1.529) inoculated cats on days 3 and 5 post-infection (pi). Note, intensive labeling of NP (Green) on the tissues from D614G-, and Delta-infected cats, while trace amount of staining on tissues from Omicron-infected animals. Nuclear counterstain was performed with DAPI (Blue) (**A**). In addition to above described tissues, palate/tonsil, retropharyngeal lymph node, and heart were evaluated. IFA scores (established as described in the Materials and Methods) for the tissues evaluated on days 3 and 5 pi are presented in (**B**) and (**C**), respectively.

In the nasal turbinate from D614G- and Delta-inoculated cats, epithelial cells of the nasal mucosa were the predominant cell type positive for the virus, with extensive hybridization observed on day 3 pi, and was slightly less abundant on day 5 pi. Additionally, middle and basilar portions of the epithelium were also sporadically stained (Fig 4A and 5A). Interestingly, in the trachea localized viral staining was detected in cells within the submucosal interstitial stroma, as well as cells associated with submucosal glandular or vascular elements in the trachea from D614G- and Delta-infected cats on days 3 pi, which was decreased by day 5 pi, with tracheal epithelial cells positive in only one D614G-inoculated cat. Similarly, in the lung, sparse staining of bronchial cells was observed in both animals on day 3 and 5 pi. Staining in the lung was more frequent in the interstitial regions especially in cells of the bronchiolar glands. In all tissues analyzed, viral RNA and N protein hybridization/staining in Omicron-inoculated cats were consistently lower than that observed in tissues from D614G- and Delta-inoculated cats (Fig 4A and 5A). The differences in RNA and N hybridization/staining are reflected by the ISH and IFA scores in each inoculated group (Fig 4B-D, Fig 5B and C). Together, these results demonstrate that the SARS-CoV-2 Omicron BA.1.1 presents limited tissue distribution, infection and replication sites compared to the SARS-CoV-2 D614G and Delta variants in cats.

### SARS-CoV-2 Omicron BA.1.1 causes limited lung inflammation in cats

Histological evaluation was performed in tissues collected on days 3 (*n* = 2 cats per virus inoculated group), 5 (*n* = 2 cats per group) and 14 pi (*n* = 3 cats per group) from D614G-, Delta-, and Omicron-inoculated cats (Fig 6). In the D614G group, animals had aggregates of fibrin, cellular debris, degenerate epithelial cells and leukocytes that partially filled the nasal passages on days 3 and 5 pi. Vasculitis was not observed in the nasal tissue. One animal euthanized on day 3 pi had epithelial necrosis and degeneration of the tracheal epithelium, while epithelial degeneration, necrosis, and regeneration associated with a mixed inflammatory infiltrate was observed in all animals on days 3, 5 and 14 pi, in which these changes were less on day 14 pi (Fig 6A-D). Examination of the lung revealed one animal with bronchial and bronchiolar necrosis and degeneration accompanied by sloughed epithelial cells and mixed inflammation on day 3 pi (Fig 6A-D). Additionally, two animals had mixed inflammation in the alveoli on day 3 and one animal had perivascular edema. No histologic lesions were noted in the palate/tonsil, retropharyngeal lymph node, and heart.

**Figure 6.**
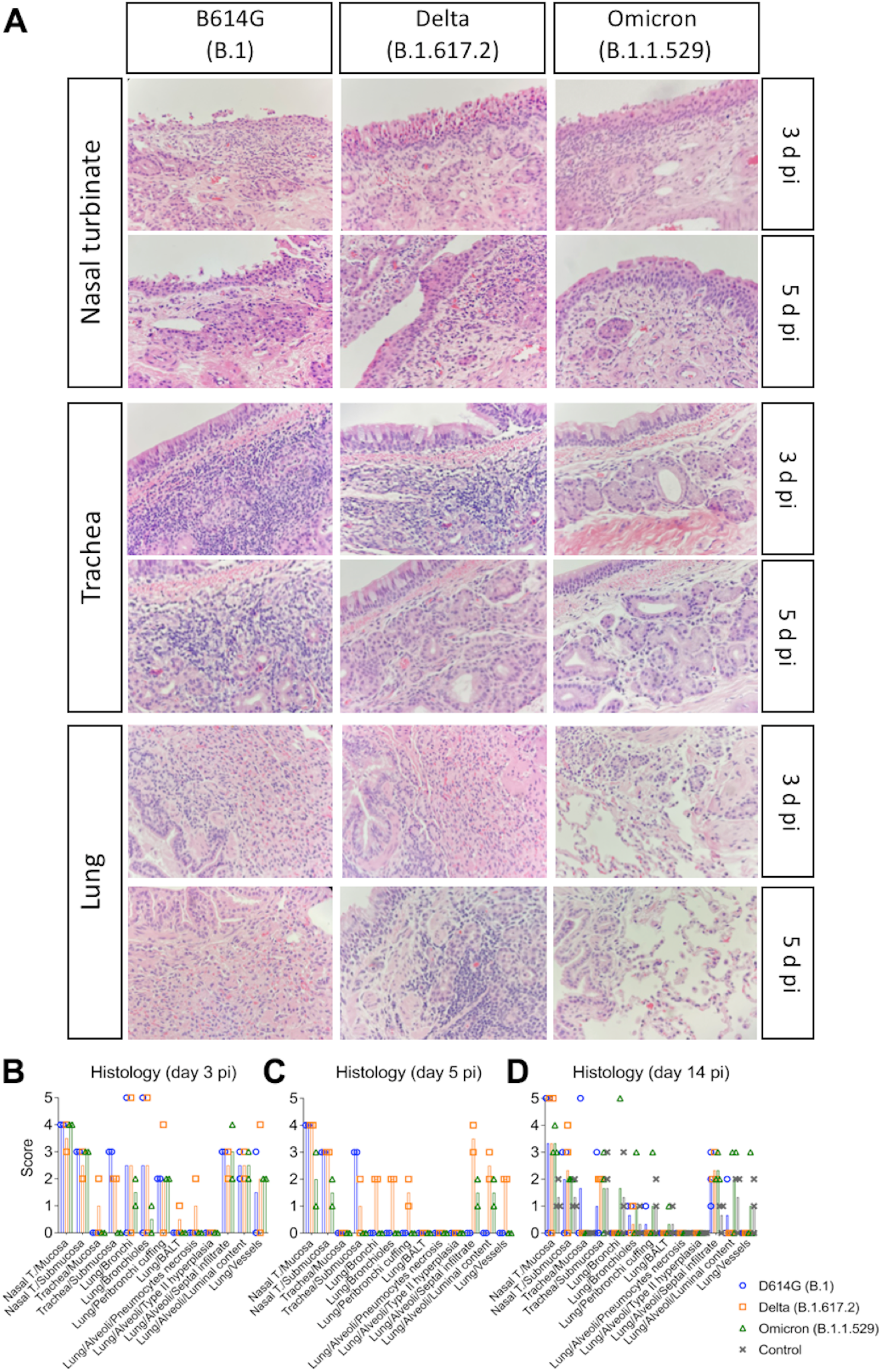
Histological examination of respiratory tract of SARS-CoV-2 D614G, Delta, and Omicron inoculated cats. Histological examination of the respiratory tract from SARS-CoV-2-inoculated cats on days 3, and 5 post-infection (pi). Intense inflammatory infiltrates in the nasal turbinate can be observed. Trachea and lung are less compromised, and the lesions in the lung of Omicron-inoculated cats are less intense (**A**). Lesion scores (established as described in the Materials and Methods) for the tissues evaluated on days 3, 5, and 14 pi are presented in (**B**), (**C**), and (**D**), respectively.

In the Delta variant inoculated group, animals presented significant epithelial necrosis, degeneration, and regeneration of the nasal epithelium accompanied by aggregates of fibrin, cellular debris, degenerate epithelial cells and leukocytes that partially filled the nasal passages mainly on days 3 and 5 pi (Fig 6A-D). One animal had fibrinoid vascular necrosis and vasculitis in the nasal passages. In the lung, marked bronchial and bronchiolar epithelial degeneration and necrosis were observed in one animal on days 3 and 5 pi, whereas the other animals did not have evidence of pulmonary necrosis. Four animals euthanized on days 3 and 5 pi had mixed inflammation in the lumina of bronchi and bronchioles. Mixed inflammation was noted in the interstitium of all animals with one animal having concurrent hyaline membranes and vasculitis of interstitial capillaries. Peribronchial and perivascular edema was present in one animal. No lesions were noted in the palate/tonsil, trachea, retropharyngeal lymph node, and heart.

Histological changes observed in the upper respiratory tract of Omicron-inoculated cats consisted of epithelial necrosis in the nasal cavity with variably epithelial attenuation and regeneration on days 3, 5 and 14 pi (Fig 6A-D). Additionally, all animals euthanized on days 3 and 5 pi and one cat euthanized on day 14 pi had large fibrin-rich aggregates admixed with sloughed epithelial cells and mixed inflammation in the nasal cavity. Three out of 7 animals had significant mixed perivascular inflammation with one animal having overt fibrinoid vascular necrosis and vasculitis. In contrast to histological features observed in D614G and Delta-inoculated cats, no necrotizing lesions were observed in the lung of any Omicron-inoculated animals (Fig 6A-D). Two animals had intraluminal mixed inflammation in the bronchioles and bronchi. All animals had mixed inflammation present in alveoli, which was associated with fibrin deposition in one of the animals. Peribronchial and perivascular edema was present in most animals. One animal had mixed inflammation present in the submucosa of the palate tissue, and no lesions were noted in the trachea, retropharyngeal lymph node, and heart.

Necrotizing lesions were not observed in the lung of control animals. Rare alveolar histiocytosis accompanied by peribronchial, peribronchiolar, and perivascular mononuclear infiltrates were present. Mild pulmonary edema was present in two animals. Two of the control animals had epithelial degeneration and necrosis in the nasal mucosa. This was accompanied by mild fibrin exudation in one animal. All animals had mild mucosal and submucosal mixed inflammation in the nasal cavity. No lesions were noted in the palate/tonsil, retropharyngeal lymph node, trachea, and heart.

Together, these results demonstrate that the SARS-CoV-2 Omicron BA.1.1 presents a limited pathogenicity in the infected cats compared to that of SARS-CoV-2 D614G and Delta variants.

### Antibody responses following SARS-CoV-2 D614G, Delta and Omicron infection in cats

The antibody responses SARS-CoV-2 infection in cats were assessed by an indirect ELISA and by plaque reduction neutralization (PRNT) and virus neutralization (VN) assays (Fig 7A-C). Using a N protein-based indirect ELISA with a defined cut-off value (S/P ratio) of 0.5, antibodies against the N protein were detected as early as day 7 pi for all three cats in the D614G variant inoculated group (Mean S/P: 0.73) and in Delta variant infected group (Mean S/P: 0.62), while only two out of three cats in Omicron variant infected group showed low seroconversion by day 7 pi (Mean S/P: 0.59). No significant difference was observed for the antibody levels between groups at day 7 pi. At day 14 pi, all cats from D614G, Delta, and Omicron inoculated groups presented a high antibody level with mean S/P ratio of 0.92, 1.72, and 0.81, respectively. Delta variant induced highest antibody response with significant differences (*p* < 0.01) when compared with the other variant viruses on day 14 pi (Fig 7A). No antibody responses were detected in control group animals.

**Figure 7.**
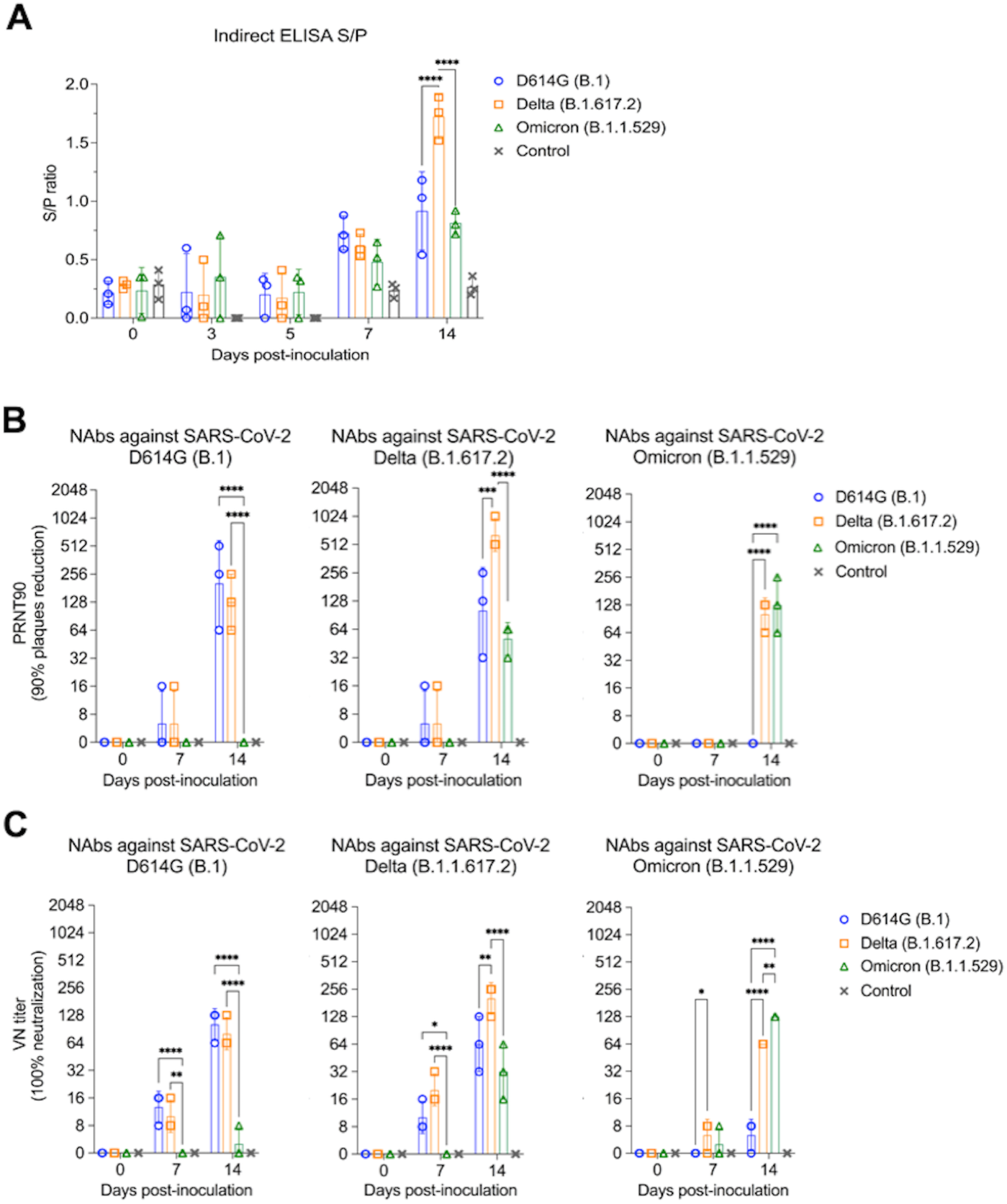
Serological responses following SARS-CoV-2 D614G, Delta, and Omicron inoculation in cats. Kinetics of antibody response against SARS-CoV-2 N protein. Serum from SARS-CoV-2 D614G (B.1), Delta (B.1.617.2), and Omicron BA.1.1 (B.1.1.529) inoculated cats collected at days 0, 3, 5, 7, and 14 pi were subjected to indirect ELISA analysis. Y-axis represents S/P ratio and different colors denotes different treatment groups. Data are presented as means ± standard error (**A**). Neutralizing antibodies (NAbs) responses to SARS-CoV-2 and cross reactivity between D614G (B.1), Delta (B.1.617.2), and Omicron BA.1.1 (B.1.1.529) variants were assessed by PRNT90 (90% plaques reduction) (**B**) and VN (100% neutralization) (**C**). Data are presented as means ± standard deviation. * *p <* 0.05; ** *p <* 0.01; *** *p <* 0.005; **** *p <* 0.0001.

Neutralizing antibodies (NAbs) responses to SARS-CoV-2 and cross reactivity between D614G, Delta, and Omicron variants were assessed by PRNT and VN assays in serum samples collected on days 0, 7, 14 pi (*n* = 3 cats per group). All D614G- and Delta-inoculated cats showed NAbs titers to SARS-CoV-2 as early as day 7 pi, while NAbs in Omicron-inoculated cats were only detected on day 14 pi (Fig 7B and C). Cross neutralization assays revealed cross-neutralizing activity of serum from D614G-inoculated cats against the Delta variant, but not against the Omicron variant. Interestingly, cross neutralizing antibodies induced by infection with the Delta variant were detected against D614G and Omicron variants. Omicron infection induced lower and delayed neutralizing antibodies, which were efficient in neutralizing both Omicron and Delta variants (Fig 7B and C). All the controls animals remained seronegative throughout the experiment. These results confirm infection of all SARS-CoV-2 D614G-, Delta- and Omicron inoculated cats, demonstrating the susceptibility of domestic cats to SARS-CoV-2 variants.

### Inflammatory cytokine response in SARS-CoV-2 infected cat serum

A multiplex Luminex bead-based immunoassay was used to measure serum cytokine/chemokine responses over the course of infection. SARS-CoV-2 D614G- and Delta-inoculated cats showed increased inflammatory cytokines (Fig 8), including a high-level production of IL-2, GM-CSF, KC at day 3 pi (Fig 8D, G and H) and IL-1β, TNF-α, IL-2, IL-4, IL-6, KC (Neutrophil chemoattractant) at day 7 pi (Fig 8 A, B, D-F, H). Interestingly, in comparison to the SARS-CoV-2 D614G- and Delta-inoculated cats, Omicron variant infection induced increased production of IFN-γ at days 7 and 14 pi (Fig 8C), higher level of RANTES (Regulated on Activation, Normal T Cell Expressed and Secreted) at days 3 and 5 pi (Fig 8I).

**Figure 8.**
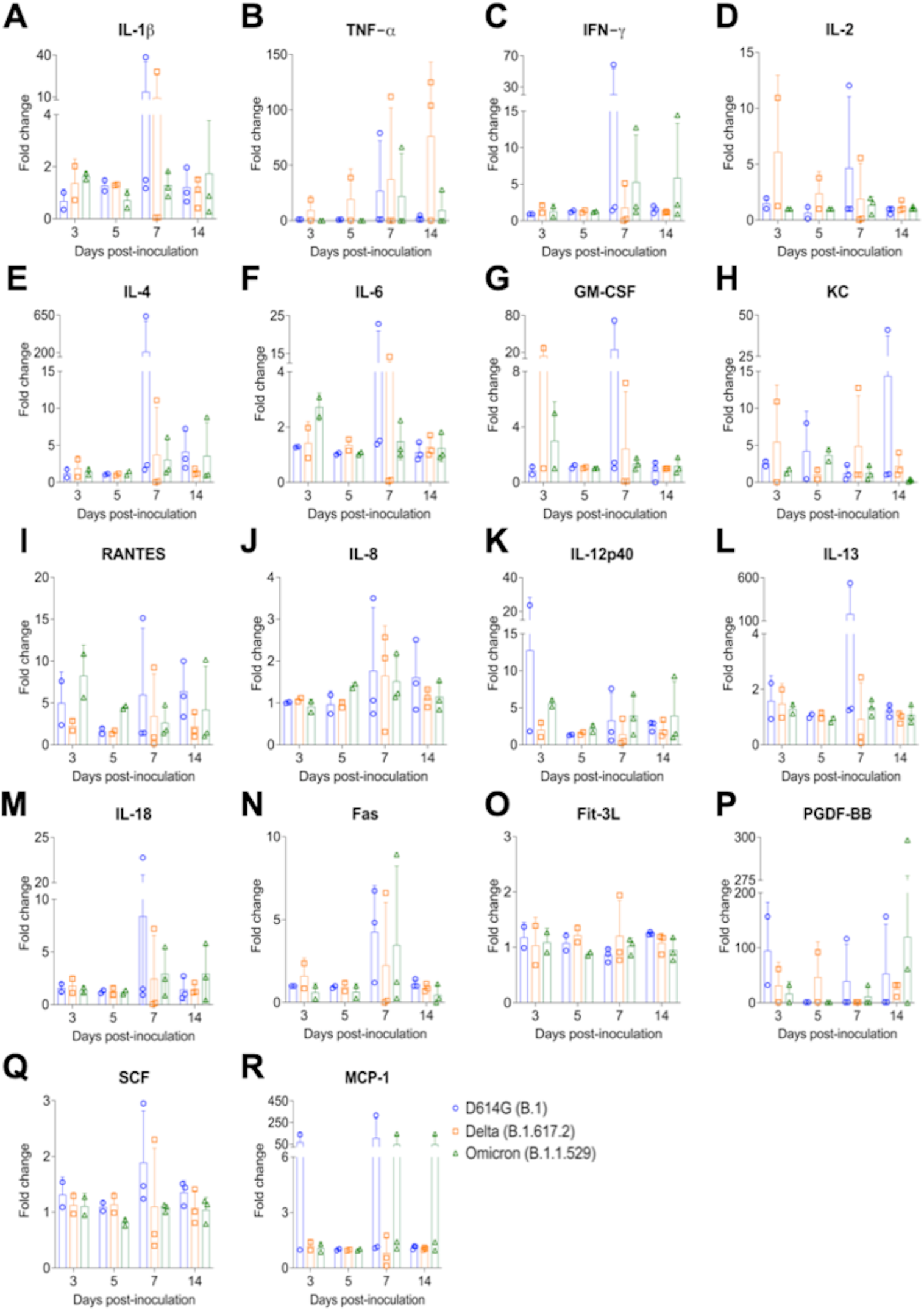
Serum inflammatory cytokine response after infection with different SARS-CoV-2 variants. **A-R**) Serum from SARS-CoV-2 D614G (B.1), Delta (B.1.617.2), and Omicron BA.1.1 (B.1.1.529) inoculated cats collected at days 3, 5, 7, and 14 pi were subjected to Milliplex™ immunoassay to determine the protein expression levels of cytokines/chemokines. X-axis represents different time points and Y-axis represents fold change relative to each corresponding cat at day 0 pi. Data are presented as means ± standard error. KC = neutrophil chemoattractant; RANTES (CCL5) = Regulated upon Activation, Normal T Cell Expressed and Secreted; TNF-α = Tissue Necrosis Factor α; GM-CSF = Granulocyte-macrophage colony-stimulating factor.

## Discussion

Since the first reported cases of SARS-CoV-2 infection in humans [20], the COVID-19 pandemic has affected nearly every country in the world. The virus remains evolving while circulating in the human population, leading to the emergence of viral variants [9]. SARS-CoV-2 spike (S) protein plays a key role in interaction with the host ACE2 receptor. Mutations in S protein have been shown to impact virus infectivity, transmissibility and virulence in humans and animal models [21,22]. Here we characterized the infection dynamics, tissue tropism and compared the pathogenesis of SARS-CoV-2 D614G (B.1), Delta (B.1.617.2) and Omicron BA.1.1 (B.1.1.529) variants in a feline model. While all three viral variants presented tropism for the respiratory tract with marked evidence of replication in the upper respiratory tract, cats inoculated with the Omicron BA.1.1 variant shed less infectious virus in respiratory secretions, and presented limited virus replication in the upper and lower respiratory tract when compared to animals inoculated with the D614G or Delta variants.

Following intranasal inoculation with each virus variant, SARS-CoV-2 D614G- and Delta-inoculated cats became lethargic with increased body temperatures on days 1 to 3 pi, whereas Omicron-inoculated cats remained subclinical for the duration of the experiment. Omicron-inoculated cats gained weight throughout the 14-day experimental period similarly to what was observed in the control (mock inoculated) animals, suggesting limited pathogenicity of the Omicron variant in the domestic cat model. Interestingly, low pathogenicity of the Omicron variant has been showed in murine models of SARS-CoV-2 infection. While SARS-CoV-2 D614G- or Delta-variant infection caused weight loss in Syrian hamsters and humanized mice, lower or no body weight losses were observed in Omicron inoculated animals [15,18,23–25].

Notably, although the emergence of the Omicron variant in the human population led an increase in the number of COVID-19 cases, mild symptoms, lower viral loads, as well as lower risk of hospitalization and death have been described in Omicron-infected people in comparison to infections with previous variants [26–31]. Similarly, in addition to the lower pathogenicity of SARS-CoV-2 Omicron in cats (Figure 9), virus replication and infectious virus shedding was significantly lower than in D614G- and Delta-inoculated animals (*p* < 0.001) (<3.1 log10 TCID_50_.ml^-1^, and up to 6.3 log10 TCID_50_.ml^-1^, respectively). Moreover, lower viral shedding in respiratory secretions was followed by reduced viral load in tissues of SARS-CoV-2 Omicron-inoculated cats. The highest viral titers were observed in the nasal turbinate (∼6.0 log10 TCID_50_.ml^-1^) from D614G- and Delta-inoculated cats on days 3 and 5 pi, while markedly lower viral loads were detected in Omicron-inoculated animals (up to 3.0 log10 TCID_50_.ml^-1^). In addition, replicating virus, measured by subgenomic viral RNA and infectious virus quantification, in the trachea and lung of Omicron-inoculated cats were lower than those observed in D614G- and Delta-inoculated animals (Figure 9). While viral loads in actual human tissues during SARS-CoV-2 infection is unknown, experimental inoculation studies in animal models have shown that tissue distribution of SARS-CoV-2 Omicron in Syrian hamsters and humanized mice is reduced when compared to the D614G and Delta variants [15,23–25].

**Figure 9.**
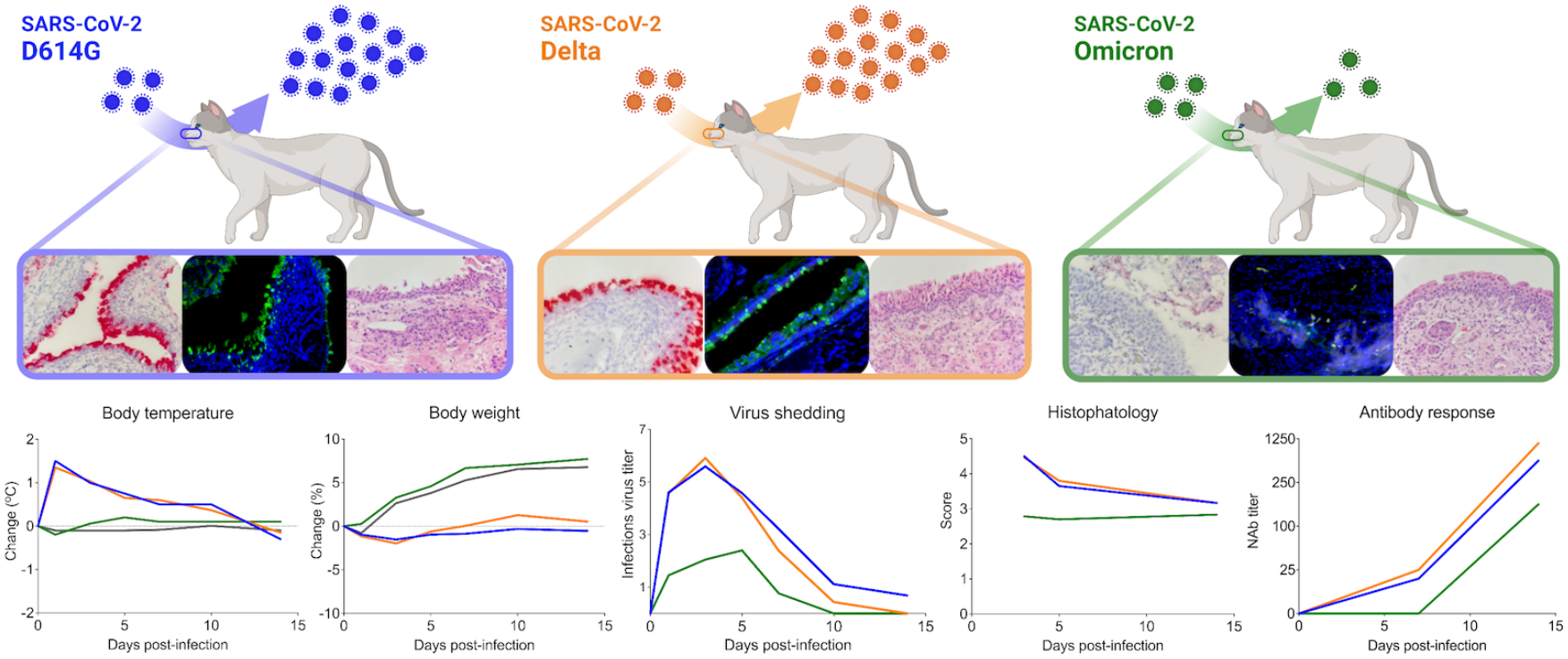
Infection dynamics and pathogenesis of SARS-CoV-2 D614G, Delta and Omicron BA.1.1 variants in a feline model of infection. After intranasal inoculation with SARS-CoV-2 D614G or Delta, cats became lethargic, and showed increased body temperatures, while Omicron-inoculated and controls cats (mock inoculate) (grey lines in the graphs) remained subclinical and gained weight throughout the experimental period. Cats inoculated with SARS-CoV-2 D614G- and the Delta variants presented in higher levels of infectious virus shedding in nasal secretions and in tissues, whereas strikingly lower levels of virus shedding and reduced tissue distribution and histologic lesions were observed on Omicron-inoculated animals. Neutralizing antibody (NAbs) responses were higher in SARS-CoV-2 D614G or Delta inoculated cats.

Consistent with SARS-CoV-2 subgenomic RNA and infectious virus quantification in tissues, the distribution of Omicron was markedly reduced in comparison to D614G and Delta variants, as evidenced lower by *in situ* viral RNA detection, immunofluorescence staining for the N protein in the upper and lower respiratory tract of inoculated cats. Importantly, tissue distribution of viral RNA and N protein reflected the degree and severity of the histological changes observed in tissues. The differences were most evident in the lungs, with Omicron-inoculated cats only presenting limited inflammatory responses when compared to D614G and Delta inoculated animals. A similar phenotype was observed in hACE2 transgenic mice and in wild-type and hACE2 transgenic hamsters that inoculated with SARS-CoV-2 Omicron variant [14,15,18,23–25]. While the three viruses inoculated in cats in this study presented a similar tissue tropism - primarily nasal turbinate, trachea, and lung - only scarce virus hybridization/staining was observed in tissues from Omicron-infected animals. In humans, the respiratory tract was described as the main site of SARS-CoV-2 infection and replication, which includes nasal and oropharyngeal tissues, trachea and lungs [32,33]. A study describing the SARS-CoV-2 distribution and cell specificity across the human body from 44 acute cases of COVID-19 demonstrated that the virus has extensive tissue distribution, however viral RNA was mainly detected in the respiratory tract [34].

Much has been explored about the role of cytokines in the pathogenesis of COVID-19 in humans. In this study, we assessed the expression levels of 18 feline cytokines over the time course of SARS-CoV-2 infection in cats. SARS-CoV-2 D614G- and Delta-variant infection induced production of inflammatory cytokines, including high-level of IL-2, GM-CSF, KC at day 3 pi and IL-1β, TNF-α, IL-2, IL-4, IL-6, KC (Neutrophil chemoattractant) at day 7 pi. These results are consistent with a previous report, in which excessive production of TNF-α, IL-6, IL-1β is associated with the disease severity [35]. Interestingly, in comparison to the other variants, SARS-CoV-2 Omicron-inoculated cats showed increased production of IFN-γ at days 7 and 14 pi, and RANTES (Regulated on Activation, Normal T Cell Expressed and Secreted) at days 3 and 5 pi. Consistent with this observation, a recent study reveals that decreased disease severity by Omicron variant may due to increased anti-inflammatory IFN-γ and decreased pro-inflammatory cytokine responses (IL-6) [36]. In the future, it would be interesting to assess and compare expression of such cytokines in tissues, particularly targeting the most severely affected tissues in the upper and lower respiratory tract of infected cats.

The Omicron variant sublineage BA.1.1 used in the present study differs from its sister BA.1 clade by a unique spike protein substitution R346K, which has been linked to immune escape [37–39]. Interestingly, while reciprocal cross neutralization was elicited by infection with D614G and Delta and by Delta and Omicron variants in cats, convalescent sera from Omicron infected cats did not efficiently neutralize D614G virus as evidenced in VN and PRNT assays. Consistent with these findings, serum from D614G inoculated animals did not efficiently neutralize the Omicron variant. These results indicate that neutralizing antibodies elicited against the Delta variant in cats have broader neutralizing activity, and efficiently neutralize both D614G and Omicron variants. As the Omicron variant continues to evolve and descendent sublineages including BA.2, BA.2.12.1, BA.3, BA.4 and BA.5 have been emerged [9], it will be important to conduct studies to assess their biological properties in animal models. Notably, a recent study demonstrated that the BA.2 sublineage presents replication and pathogenicity comparable to that of BA.1 in a mouse and a hamster model [40]. As a species that is naturally susceptible to SARS-CoV-2 infection [41–43] and that has been shown to efficiently transmit the virus [44–46], domestic cats represent an invaluable animal model to assess the infectivity and pathogenicity of SARS-CoV-2 variants.

Understanding the infectivity and pathogenesis of SARS-CoV-2 VOCs is essential to develop improved vaccines and therapeutics to effectively control the COVID-19 pandemic. Here we showed that while D614G- and Delta-inoculated cats presented severe pneumonia, histopathological examination of the lungs from Omicron-infected cats revealed absence or only low focal inflammation, which also correlated to the significant differences in viral loads, replication properties in other tissue sites and host immune responses. Together, these results (as summarized in Fig 9) demonstrate that the Omicron BA.1.1 variant is less pathogenic than the D614G and Delta variants in a highly susceptible feline model of SARS-CoV-2 infection.

## Methods

### Cells and viruses

Vero E6 (ATCC^®^ CRL-1586™), and Vero E6/TMPRSS2 (JCRB Cell Bank, JCRB1819) were cultured in Dulbecco’s modified eagle medium (DMEM), supplemented with 10% fetal bovine serum (FBS), L-glutamine (2mM), penicillin (100 U.ml^−1^), streptomycin (100 μg.ml^−1^) and gentamycin (50 μg.ml^−1^). The cell cultures were maintained at 37 °C with 5% CO_2_. The SARS-CoV-2 isolates used in this study were obtained from residual human anterior nares, or nasopharyngeal secretions. The SARS-CoV-2 D614G (B.1 lineage) New York-Ithaca 67-20 (NYI67-20), and Delta (B.1.617.2 lineage) NYI31-21 isolates, were propagated in Vero E6/TMPRSS2 cells, whereas the Omicron BA.1.1 (B.1.1.529) NYI45-21 isolate was propagated in Vero E6 cells. Low passage virus stock (passage 3) were prepared, cleared by centrifugation (2000 x *g* for 15 min) and stored at −80 °C. The whole genome sequences of the virus stocks were determined to confirm that no mutations occurred during amplification in cell culture. The titers of virus stock were determined by plaque assays, calculated according to the Spearman and Karber method and expressed as plaque-forming units per milliliter (PFU.ml^−1^).

### Ethics statement

Viral isolates were obtained from residual de-identified human anterior nares or nasopharyngeal secretions collected as part of the Cornell University surveillance program or at the Cayuga Medical Center Inovation laboratory. The protocols and procedures were reviewed and approved by the Cornell University and Cayuga Medical Center Institutional Review Boards (IRB approval numbers 2101010049 and 0420EP, respectively). All animals were handled in accordance with the Animal Welfare Act. The study procedures were reviewed and approved by the Institutional Animal Care and Use Committee at the Cornell University (IACUC approval number 2020-0064).

### Animals housing and experimental design

A total of twenty-four 24-40-month-old domestic cats (*Felis catus*) (four males and three females [*n* = 7] per inoculated group, and two males and one female [*n* = 3] for the control group) were obtained from Clinvet (Waverly, NY, USA). Animals were donated to Cornell University to support the reduction of animal use in research. All animals were housed in the animal biosafety level 3 (ABSL-3) facility at the East Campus Research Facility (ECRF) at Cornell University. After acclimation, cats were anesthetized and inoculated intranasally with 1 ml (0.5 ml per nostril) of a virus suspension containing 5 x 10^5^ PFU of SARS-CoV-2 D614G (B.1 lineage), Delta (B.1.617.2 lineage), or the Omicron BA.1.1 (B.1.1.529) variants. Control cats were mock-inoculated with Vero E6/TMPRSS2 cell culture medium supernatant. All animals were maintained individually in Horsfall HEPA-filtered cages, connected to the ABSL-3’s exhaust system. Body temperatures and weight were measured on a daily basis. Oropharyngeal (OPS), nasal (NS), and rectal swabs (RS) were collected under sedation (dexmedetomidine) on days 0, 1, 3, 5, 7, 10, and 14 post-inoculation (pi). Upon collection, swabs were placed in sterile tubes containing 1 ml of viral transport medium (VTM Corning®, Glendale, AZ, USA) and stored at −80 °C until processed for further analyses. Blood was collected under sedation (dexmedetomidine) through jugular venipuncture using a 3 ml sterile syringe and 21G x 1” needle and transferred into serum separator tubes on days 0, 3, 5, 7, 10 and 14 pi. The blood tubes were centrifuged at 1200 x *g* for 10 min and serum was aliquoted and stored at −20 °C until further analysis. Cats were humanely euthanized on day 14 pi. Following necropsy, tissues, including nasal turbinate, palate/tonsil, retropharyngeal LN, trachea, lung, mediastinal LN, heart, liver, spleen, kidney, small intestine, mesenteric LN were collected and processed for rRT-PCR and virus titration. Additionally, tissue samples were collected and processed for standard microscopic examination, a subset of tissues were also processed by *in situ* hybridization (ISH) and *in situ* immunofluorescence (IFA). For this, tissue sections of approximately 0.5 cm in width were fixed by immersion in 10% neutral buffered formalin (20 volumes fixative to 1 volume tissue) for approximately 72 h, and then transferred to 70% ethanol, followed by standard paraffin embedding techniques. Slides for standard microscopic examination were stained with hematoxylin and eosin (HE).

### Nucleic acid isolation and real-time reverse transcriptase PCR

Nucleic acid was extracted from NS, OPS, RS, BALF and tissue samples collected at necropsy. For NS, OPS, RS, and BALF samples 200 µL of cleared swab supernatant were used for nucleic acid extraction. For tissues, 0.3 g of each tissue were minced with a sterile disposable scalpel, resuspended in 3 ml DMEM (10% w/v) and homogenized using a stomacher (one speed cycle of 60s, Stomacher® 80 Biomaster). Then, the tissues homogenate supernatant were centrifuged at 2000 x *g* for 10 min and 200 µL of cleared supernatant was used for RNA extraction using the MagMax Core extraction kit (Thermo Fisher, Waltham, MA, USA) and the automated KingFisher Flex nucleic acid extractor (Thermo Fisher, Waltham, MA, USA) following the manufacturer’s recommendations. The real-time reverse transcriptase PCR (rRT-PCR) for total viral RNA detection was performed using the EZ-SARS-CoV-2 Real-Time RT-PCR assay (Tetracore Inc., Rockville, MD, USA), which detects both genomic and subgenomic viral RNA targeting the virus nucleoprotein (N) gene. An internal inhibition control was included in all reactions. Positive and negative amplification controls were run side-by-side with test samples. For specific subgenomic RNA detection, a RT-qPCR reaction targeting the virus envelope protein (E) gene was used following the primers and protocols previously described [19]. Both RT-PCR (for total viral RNA detection) and RT-qPCR assay (for specific subgenomic RNA detection) were validated using a standard curve by using ten-fold serial dilutions from 10^0^ to 10^−8^ of virus suspension containing 10^6^ TCID_50_.ml^-1^ for each of the SARS-CoV-2 variants used in the study. Relative viral genome copy numbers were calculated based on the standard curve and determined using GraphPad Prism 9 (GraphPad, La Jolla, CA, USA). The amount of viral RNA detected in samples were expressed as log10 (genome copy number) per ml.

### Virus isolation and titrations

Samples that tested positive for SARS-CoV-2 by rRT-PCR were subjected to virus isolation under Biosafety Level 3 (BSL-3) conditions at the Animal Health Diagnostic Center (ADHC) Research Suite at Cornell University. NS, OPS, RS, BALF, and tissues homogenate supernatant were subjected to end point titrations. For this, samples were subjected to limiting dilutions and inoculated into Vero E6/TMPRSS2 cells cultures prepared 24 h in advance in 96-well plates. At 48 h post-inoculation, cells were fixed and subjected to an immunofluorescence assay (IFA) as described in a previous study [47]. Virus titers were determined on each time point using end-point dilutions and the Spearman and Karber’s method and expressed as TCID_50_.ml^−1^.

### *In situ* RNA detection

Paraffin-embedded tissues from days 3, 5, and 14 pi were sectioned at 5 µm and subjected to *in situ* hybridization (ISH) using the RNAscope^®^ ZZ probe technology (Advanced Cell Diagnostics, Newark, CA). Tissues from inoculated and controls cats including nasal turbinate, palate/tonsil, retropharyngeal lymph nodes, trachea, lung, and heart were subjected to ISH using the RNAscope^®^ 2.5 HD Reagents–RED kit (Advanced Cell Diagnostics) following the manufacturer’s instructions, and using a probe targeting SARS-CoV-2 RNA spike (V-nCoV2019-S probe ref # 848561). A probe targeting feline host protein peptidylprolyl isomerase B (PPIB) was used as a positive control (Advanced Cell Diagnostics cat # 455011). A probe targeting DapB gene from Bacillus subtilis strain SMY was used as a negative control (Advanced Cell Diagnostics cat # 310043). Tissue sections were ISH scored based on labeling extension using a scoring system as follow: score 1 = up to 2% of the tissue section positive; score 2 = from 2 to 5% positive; score 3 = from 5 to 15% positive; score 4 = from 15 to 25%; score 5 = more than 25%; and score 0 = no labeling. Vero E6 cell infected with 0.1 MOI of SARS-CoV-2 D614G (B.1 lineage), Delta (B.1.617.2 lineage), or the Omicron BA.1.1 (B.1.1.529) and fixed at 24 h pi were used as positive controls (Suppl. Fig 1).

### *In situ* immunofluorescence

Paraffin-embedded tissues from days 3 and 5 pi were sectioned at 5 µm and subjected to immunofluorescence assay (IFA). Tissues from inoculated and control cats including nasal turbinate, palate/tonsil, retropharyngeal lymph nodes, trachea, lung, and heart were subjected to IFA. Formalin-fixed paraffin-embedded (FFPE) tissues were deparaffinized with xylene and rehydrated through a series of graded alcohol solutions. Antigen unmasking was performed using Tris-based antigen unmasking solution pH 9.0 (Vector Laboratories ref # H-3301) by boiling the slides in the unmasking solution for 20 min. After 10 min at 0.2% Triton X-100 (in phosphate-buffered saline [PBS]) at room temperature (RT), and 30 min blocking using a goat normal serum (1% in PBS) at RT, tissues were subjected to IFA. A mouse monoclonal antibody targeting nucleoprotein (N) of SARS-CoV-2 was used as a primary antibody (SARS-CoV-2 N mAb clone B61G11) [48,49] was incubated for 45 min at RT. Followed by 30 min incubation at RT with a goat anti-mouse IgG antibody (goat anti-mouse IgG, Alexa Fluor® 488). Nuclear counterstain was performed with 4’,6-Diamidino-2-Phenylindole, Dihydrochloride (DAPI) (10 min at RT). Tissue sections were IFA scored based on staining extension using scoring system as described above. Vero E6 cell infected with 0.1 MOI of SARS-CoV-2 D614G (B.1 lineage), Delta (B.1.617.2 lineage), or the Omicron BA.1.1 (B.1.1.529) viruses were fixed at 24 h pi and used as positives controls (Suppl. Fig 2).

### Histology

For the histological examination, tissue sections of approximately 0.5 cm in width were fixed by immersion in 10% neutral buffered formalin (≥20 volumes fixative to 1 volume tissue) for approximately 72 h, and then transferred to 70% ethanol, followed by standard paraffin embedding techniques. Tissues collected from inoculated and controls cats including nasal turbinate, palate/tonsil, retropharyngeal lymph nodes, trachea, lung, and heart were subjected to histological examination after stained with hematoxylin and eosin (HE). Histologic examination was performed by a board certified veterinary anatomic pathologist (ADM) and histological changes were scored based on severity using a previously described scoring system [50]. For lung, bronchi, bronchioles, alveoli, blood vessels, and pleura were all analyzed for lesions based on a combination of a graded scoring system and the presence/absence of lesions. The tracheal and nasal turbinate mucosa, submucosa, and associated vessels were all scored independently. Other organs including palate/tonsil, retropharyngeal lymph node, and heart were analyzed only for the presence/absence of histologic lesions.

### Indirect Enzyme-linked immunosorbent assay (ELISA)

Indirect ELISA was developed in-house based on the modified method described previously [51,52]. The SARS-CoV-2 nucleoprotein (N) from original strain (Wuhan-hu-1) was expressed in the *E.coli* BL21 cells and purified using Ni-NTA system as described in a previous study [53]. Immulon 2HB plate (Thermo Fisher Scientific, Waltham, MA, USA) was coated with 100 μl N antigen at 300 ng/well. After incubation at 4 °C overnight, plates were washed three times with PBS containing 0.05% Tween 20 (PBST) and blocked with 5% (wt/vol) powdered dry milk in PBST for 1 hour at 37 °C. Serum samples collected at day 0, 3, 5, 7, and 14 pi were diluted 1:400 in blocking buffer and 100 µL were applied to each well. After incubation at 37 °C for 1 hour, plates were washed three times and further incubated with 100 µL horseradish peroxidase (HRP)- conjugated Goat anti-Feline IgG (H+L) secondary Antibody (Thermo Fisher Scientific, Waltham, MA, USA). The plates were incubated for 1 hour at 37 °C, and 100 µL of ABTS peroxidase substrate (KPL, Gaithersburg, MD) was added for color development followed by addition of 100 µL of ABTS stop solution (KPL, Gaithersburg, MA) for 30 min. Plates were read at 405 nm with a SpectraMax® iD5 microplate reader (Molecular Devices, San Jose, CA, USA). Sample-to-positive (S/P) ratio was calculated using the following formula: S/P = (OD of sample - OD of buffer)/(OD of positive control - OD of buffer).

### Neutralizing antibodies

Neutralizing antibody responses to SARS-CoV-2 were assessed by plaque reduction neutralization tests (PRNT) and virus neutralization (VN) assay performed under BSL-3 laboratory conditions. Serum samples collected at day 0, 7, and 14 pi were tested against the three SARS-CoV-2 variants D614G, Delta and Omicron. For the PRNT assay, twofold serial dilutions (1:16 to 1:1,024) of serum samples were incubated with 50 - 100 PFU of SARS-CoV-2 D614G, Delta, or Omicron viruses for 1 h at 37 °C. Following incubation, mixture of serum and virus were transferred to a Vero E6 cell monolayer prepared 24 h in advance in a 6-well plate and incubated for 1 h at 37 °C with 5% CO_2_. Following incubation, the mixture serum and virus was removed and cells overlayed with DMEM 10% FBS containing 0.5% agarose and incubated for 72 h at 37°C with 5% CO_2_. To visualize plaques, 250 µL of neutral red staining solution (0.033% diluted in DMEM 10% FBS) was added per well and incubated overnight at 37 °C with 5% CO_2_. After incubation, plaques were counted and the PRNT90 titer was determined. End point titers were considered the reciprocal of the serum dilution able to reduce et least in 90% the number of plaques in comparison to back titration.

For the VN assay, twofold serial dilutions (1:8 to 1:1,024) of serum samples were incubated with 100 - 200 TCID_50_ of SARS-CoV-2 D614G, Delta, or Omicron viruses, for 1 h at 37 °C. Following incubation of serum and virus, 50 µl of a cell suspension of Vero E6 cells was added to each well of a 96-well plate and incubated for 48 h at 37 °C with 5% CO_2_. At 48 h post-inoculation, cells were fixed and subjected to IFA as described in a previous study [47]. Neutralizing antibody titers were expressed as the reciprocal of the highest dilution of serum that completely inhibited SARS-CoV-2 infection/replication. Fetal bovine serum (FBS) and positive and negative serum samples from white-tailed-deer [47] were used as controls.

### Luminex assay for feline inflammatory cytokines/chemokines analysis

Serum samples from cats at days 0, 3, 5, 7, and 14 pi were analyzed using the MILLIPLEX® Feline Cytokine/Chemokine Magnetic Bead Panel (MilliporeSigma, Burlington, MA, USA) following the manufacturer’s instructions. Briefly, 10 μl of serum sample was mixed with matrix solution and assay buffer, and then incubated overnight with Luminex beads that immobilized with antibodies against a panel of cytokines/chemokines, including Fas, Flt-3L, GM-CSF, IFN-γ, IL-1β, IL-2, IL-4, IL-6, IL-8, IL-12 (p40), IL-13, IL-18, KC, MCP-1, PDGF-BB, RANTES, SCF, SDF-1, and TNF-α. Beads were then washed twice and incubated with 25 μl detection antibodies for 1 hour at room temperature followed by directly adding 25 μl streptavidin-phycoerythrin and incubated for another 30 min. Beads were washed for three times and analyzed using a Luminex® 200™instrument (Luminex Corp., TX, USA). Mean fluorescence intensity (MFI) was determined for each sample, followed by absolute quantification based on standard curve of each analyte calculated using Belysa® Analysis Software Version 1.2 (MilliporeSigma, Burlington, MA, USA). Results were interpreted as fold change relative to the data from each corresponding cat serum at day 0 pi and graphed using GraphPad Prism 9 (GraphPad, La Jolla, CA, USA).

### Statistical analysis and data plotting

Statistical analysis was performed by 2-way analysis of variance (ANOVA) followed by multiple comparisons and. Statistical analysis and data plotting were performed using the GraphPad Prism software (version 9.0.1). Figures 1 and 9 were created with BioRender.com.

## Acknowledgments

We thank the Center for Animal Resources and Education (CARE) staff and Cornell Biosafety team for the support. The authors also thank Dr. Cara Mitchel and all staff at Clinvet for their support and donation of the animals in an effort to reduce the animal use (3R’s). Extraction and real-time PCR testing was conducted at the Animal Health Diagnostic Center at Cornell University. This work was funded by the National Institute of Health (NIH) and National Institute of Allergy and Infectious Diseases (NIAID) (grant no. R01AI166791-01).

---

**Suppl. Fig.1.**
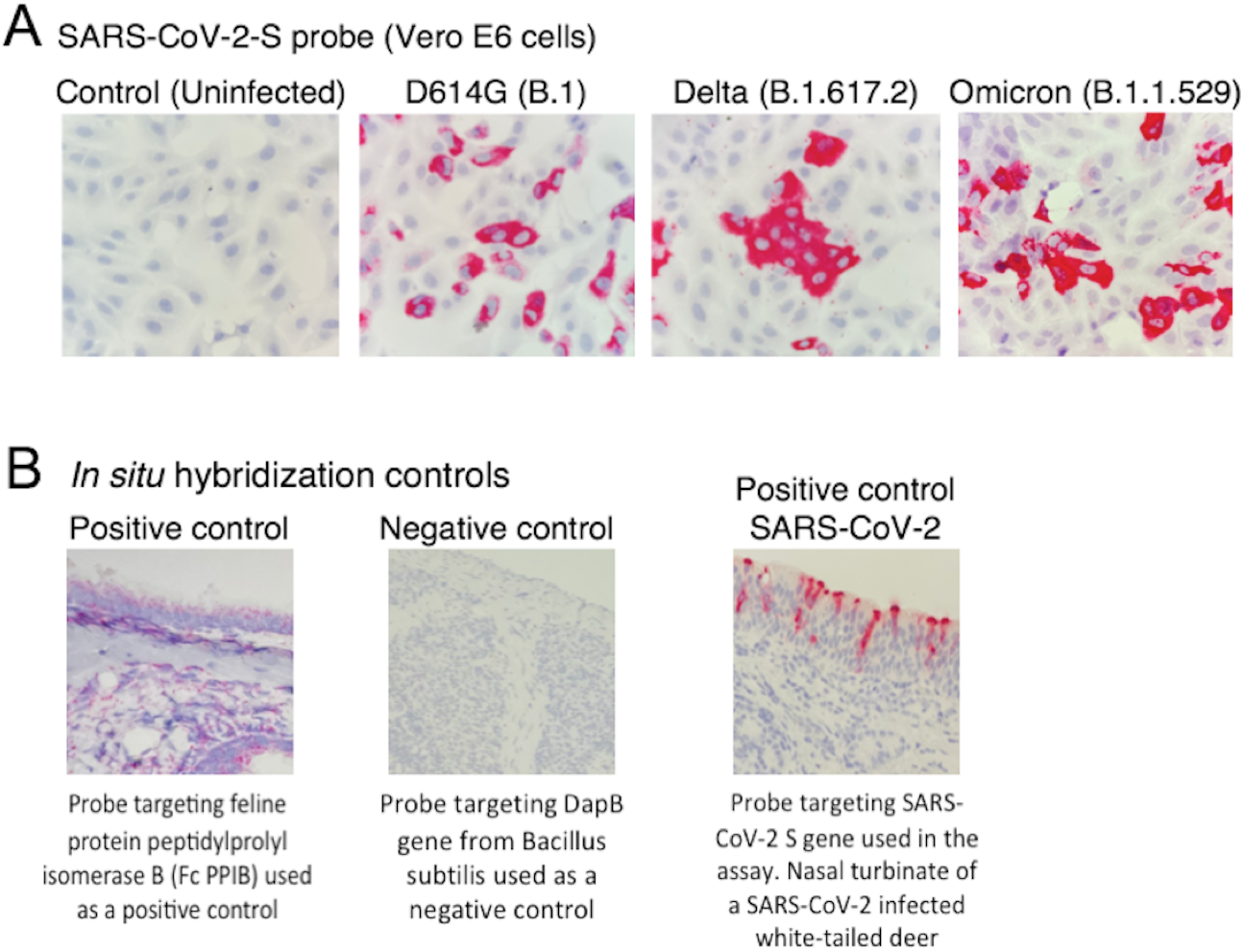
*In situ* hybridization (ISH) controls. Vero E6 cells were inoculated with SARS-CoV-2 D614G (B.1 lineage) (isolate NYI67-20), Delta (B.1.617.2 lineage) (isolate NYI31-21), and the Omicron BA.1.1 (B.1.1.529) (isolate NYI45-21) at a multiplicity of infection (MOI) of 0.1. Cells were fixed 24 h post-infection (h pi) and subjected to subjected to ISH using the RNAscope^®^ 2.5 HD Reagents–RED kit (Advanced Cell Diagnostics) following the manufacturer’s instructions using a probe targeting SARS-CoV-2 RNA spike (V-nCoV2019-S probe ref # 848561). Slides were counterstained with hematoxylin; 40x magnification (**A**). ISH using the RNAscope^®^ 2.5 HD Reagents–RED kit (Advanced Cell Diagnostics) following the manufacturer’s instructions. Left panel, nasal turbinate of a cat using a probe targeting feline host protein peptidylprolyl isomerase B (PPIB) was used as a positive control (Advanced Cell Diagnostics cat # 455011). Middle panel, nasal turbinate using a probe targeting DapB gene from Bacillus subtilis strain SMY was used as a negative control (Advanced Cell Diagnostics cat # 310043). Right panel, nasal turbinate of white-tailed deer infected with SARS-CoV-2 B.1 D614G using a probe targeting SARS-CoV-2 RNA spike (V-nCoV2019-S probe ref # 848561) was used as positive control. Slides were counterstained with hematoxylin; 40x magnification (**B**).

**Suppl. Fig.2.**
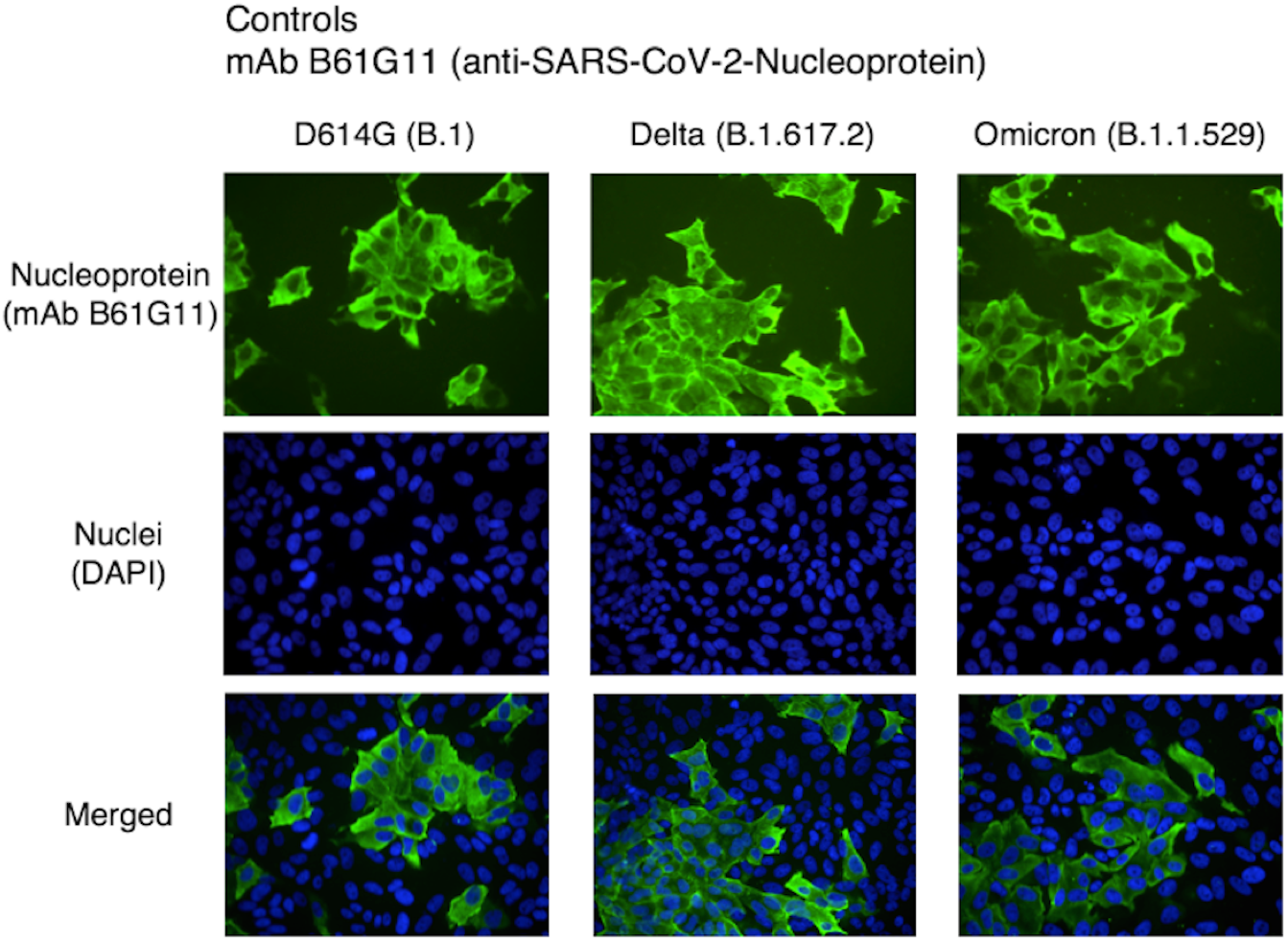
*In situ* immunofluorescence (IFA) controls. Vero E6 cell were inoculated with SARS-CoV-2 D614G (B.1 lineage) (isolate NYI67-20), Delta (B.1.617.2 lineage) (isolate NYI31-21), and the Omicron BA.1.1 (B.1.1.529) (isolate NYI45-21) at a MOI of 0.1. Cells were fixed 24 h pi and subjected to an immunofluorescence assay using a monoclonal antibody (B61G11) anti-SARS-CoV-2-nucleoprotein (N) (Green). Nuclear counterstain was performed with DAPI (Blue); 40x magnification.

